# Temporal Transcriptomic Profiling of the Developing *Xenopus laevis* Eye

**DOI:** 10.1101/2024.07.20.603187

**Authors:** Samantha J. Hack, Juli Petereit, Kelly Ai-Sun Tseng

**Affiliations:** Department of Biological Sciences, Western Michigan University, Kalamazoo, MI 49008, USA; Nevada Bioinformatics Center, University of Nevada, Reno; School of Life Sciences, University of Nevada, Las Vegas

**Keywords:** eye development, retina, *Xenopus*, ocular transcriptome, retinal progenitor cells, cell signaling, Notch

## Abstract

Retinal progenitor cells (RPCs) are a multipotent and highly proliferative population that give rise to all retinal cell types during organogenesis. Defining their molecular signature is a key step towards identifying suitable approaches to treat visual impairments. Here, we performed RNA-sequencing of whole eyes from *Xenopus* at three embryonic stages and used differential expression analysis to define the transcriptomic profiles of optic tissues containing proliferating and differentiating RPCs during retinogenesis. Gene Ontology and KEGG pathway analyses showed that genes associated with developmental pathways (including Wnt and Hedgehog signaling) were upregulated during the period of active RPC proliferation in early retinal development (Nieuwkoop Faber st. 24 and 27). Developing eyes had dynamic expression profiles and shifted to enrichment for metabolic processes and phototransduction during RPC progeny specification and differentiation (st. 35). Furthermore, conserved adult eye regeneration genes were also expressed during early retinal development including *sox2*, *pax6*, *nrl*, and Notch signaling components. The eye transcriptomic profiles presented here span RPC proliferation to retinogenesis and included regrowth-competent stages. Thus, our dataset provides a rich resource to uncover molecular regulators of RPC activity and will allow future studies to address regulators of RPC proliferation during eye repair and regrowth.

## Introduction

The incidence of preventable childhood blindness is falling, however congenital causes of severe vision loss or impairment remain due to lack of treatment options. Of specific therapeutic interest are retinal progenitor cells (RPCs), which are a multipotent population that gives rise to all adult retinal cell types including the retinal pigmented epithelium, photoreceptors, retinal ganglion cells as well as the Müller glia [1–3]. A first step towards generating new cell therapy and pharmacological treatment options is defining the molecular profile of developing optic tissues that contain cycling RPCs with the capacity to differentiate into appropriate cell types.

The African clawed frog, *Xenopus laevis* (*X. laevis*), is a powerful vertebrate system that has been used successfully for identifying the mechanisms that control normal and aberrant corneal, lens, and retinal development that lead to visual impairments [4–6]. *X. laevis* is dually advantageous as a model system because it can also regenerate a variety of mature nervous tissues including the retina, optic nerve, brain, and spinal cord [7]. It has recently been appreciated that mid-tailbud stage embryos are regrowth-competent and possess the ability to reform fully functional eyes following ablation [8,9]. Thus, *X. laevis* can be simultaneously used to investigate the similarities between developmental and regenerative transcriptional programs, as well as for identifying conserved vertebrate regulators of eye field development. This knowledge could allow researchers to address a key question of whether or how regeneration recapitulates development – an important step in providing a solid basis in therapeutic designs for visual defects and injuries.

The *X. laevis* eye field is first specified in the anterior neural plate at st. 12.5/13 [10], where the overlapping expression of eye field transcription factors (EFTFs) such as *pax6* in the eye primordium [11] specifies the RPCs committed to forming retinal tissues [12]. Retinal differentiation begins at developmental stage (st.) 24 and is completed by st. 42, over a period of two days [3,13]. During early retinal development at st. 24, RPCs are mitotic and less than 5% of RPCs have exited the cell cycle. By the mid to late retinal development periods (st. 29-38), RPCs that give rise to the presumptive ganglion cell layer (GCL) and ciliary marginal zone (CMZ) actively undergo differentiation [14–17]. Approximately 95% of RPCs exit the cell cycle by st. 37/38 [3], where progeny then continue to differentiate into the retinal cell types present in mature optic tissues. The developing lens arises from the invaginating lens placode, where the lens rudiment forms by st. 27 [4]. By st. 32, tissues become polarized into anterior and posterior domains, giving rise to the lens vesicle. Lens development is completed by st. 48 during the premetamorphic tadpole period [4].

In *Xenopus*, early embryonic stages that include specification of optic tissues have been extensively profiled by single cell RNA-sequencing, bulk RNA-sequencing, or microarray experiments [18–22]. Larval and adult stages that include the mature eye have also been profiled [18]. These studies provide a rich resource for defining the molecular details of ocular tissue specification during embryogenesis and/or characterizing gene expression profiles of the mature eye. Additionally, pathways involved in retinal fate specification are well characterized [17,23–30]. However, retinal development begins in the early tailbud embryo at st. 24 [13]. As this key temporal window has not been characterized for *X. laevis* eye development, the specific molecular drivers of RPC proliferation and differentiation in developmental contexts remain unclear.

In this study, we utilize RNA-sequencing to transcriptionally profile developing *X. laevis* eyes at three timepoints: from early tailbud stage embryos (st. 24, the beginning of retinal development) – where approximately 95% RPCs are mitotic [31], at the mid-tailbud stage (st. 27), where RPCs are regrowth-competent and remain highly proliferative, and during late retinal development (st. 35) – where ∼90% of RPCs have exited the cell cycle, to generate a mineable resource to identify regulators of RPC activity and retinogenesis. Through differential expression (DE), gene ontology (GO), and KEGG pathway analyses, we show that the transcriptional environment is primed for retinogenesis between st. 24 and 27, where markers of early RPC proliferation are more highly expressed. Our analyses identify the st. 27 eye as highly dynamic; the gene expression profile exhibits characteristics and patterns observed at st, 24 but expression begins to shift from developmental signaling programs to enrichment for molecular programs associated with phototransduction and light perception. Moreover, we found that the Notch signaling pathway is highly enriched in st. 27 optic tissues, which is a regrowth-competent stage of eye development. Notch signaling components were among many conserved eye regeneration genes expressed in developing eyes across retinogenesis, further supporting a role for Notch signaling in regulation of embryonic eye regrowth. Intriguingly, our transcriptional analysis also identified differential expression in long versus short gene homeologs across retinal development, suggesting that a detailed DE analysis examining all gene homeologs can identify novel roles for short versus long genes at different stages of embryogenesis.

## Results and Discussion

### Transcriptional Analysis of Developing Optic Tissues

To investigate the transcriptional changes in developing eyes across retinogenesis, we isolated optic tissues from *X. laevis* embryos at Nieuwkoop and Faber (NF) stages (st.) 24, 27, and 35 (Fig. 1A) and conducted bulk RNA-sequencing on poly-A enriched mRNAs. Our transcriptional dataset spans from the early tailbud stage (st. 24 and 27), when specification of retinal tissues begins in the developing optic vesicle, through retinal pigmented epithelium (RPE) morphogenesis and formation of differentiated visual structures at st. 35, a late tailbud stage (Fig. 1B).

**Figure 1:**
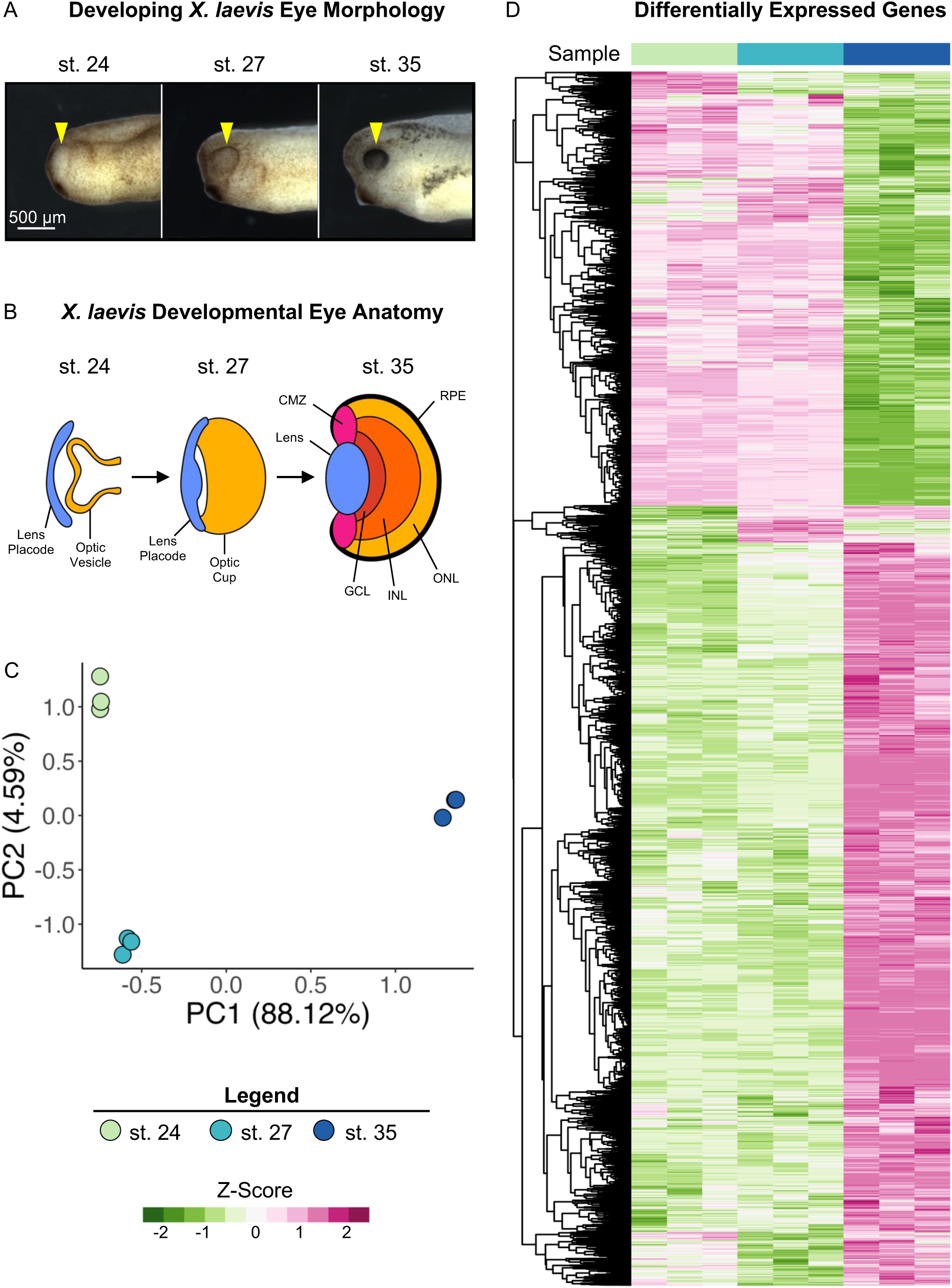
Transcriptional Analysis of Developing Optic Tissues. (**A**) *X. laevis* embryos at NF stages (st.) 24, 27, and 35 indicating the tissue that was resected for RNA-sequencing and transcriptional analysis (yellow arrowhead). Scale bar: 500 μm. **(B)** Major optic structures present at each developmental stage assayed by RNA-sequencing. CMZ: Ciliary marginal zone, GCL: Ganglion cell layer, INL: Inner nuclear layer, ONL: Outer nuclear layer, RPE: Retinal pigmented epithelium. **(C)** Principal component analysis of whole transcriptome expression in triplicate samples for each developmental stage assayed. PC1: 88.12%, PC2: 4.95. **(D)** Relative expression (Z-score vst counts) of statistically significantly differentially expressed genes between three developmental stages (st. 27 v 24 and st. 35 v 27). Rows are ordered by hierarchical clustering. Columns are ordered by stage.

Two major aligners, STAR and HISAT2 were used, and STAR showed a higher alignment rate than HISAT2 by assigning 200k to 900k more reads per sample. The overall assigned reads ranged from 19.9 to 25.6 million reads per sample. After the low count filter, the retained 17,693 transcripts were analyzed with the DESeq2 package. The principal component analysis (PCA) of the variance stabilizing transformed (vst) counts showed that developmental stages clustered in distinct groups. Transcriptional changes can be observed between all stages, with notable larger changes evidenced between st. 27 and 35 (Fig. 1C).

We conducted differential expression analysis between the three developmental stages (st. 27 vs st. 24 and st. 35 vs st. 27) to identify specific patterns of differentially expressed genes (DEGs) over the period spanning early to late retinal development, as well as transcriptional similarities. Genes with an absolute value of log_2_ fold change ≥ 1 and FDR adj. *p* < 0.05 were deemed to be statistically significantly differentially expressed (Supplemental Table 1). We identified 1,126 (427 downregulated and 699 upregulated) and 8,403 (3,074 downregulated and 5,329 upregulated) DEGs comparing st. 27 to 24 and st. 35 to 27, respectively. The PCA and differential expression analyses were well-correlated as expected; we predicted larger transcriptional changes in the later time point due to increased RPC differentiation and mature tissue formation. (Fig. 1A and 1D).

### Eye Development Gene Expression Across Retinogenesis

*X. laevis* is a well-established model of eye development, and numerous studies have described genes that have known mutant eye phenotypes. To determine if any of these known mutant genes were expressed in our transcriptional analyses, we mined the community resource *Xenbase* [32–34] for genes that produce phenotypes in the following categories when expression is manipulated: “Absence of eye,” “Abnormal development of eye,” “Decreased size of normal eye,” and “Decreased size of eye primordium” (Supplemental Table 2). We queried the expression of 70 transcripts, and only 47 homeologs (67%) had detectable levels of expression after filtering for quantifiable counts (Fig. 2A). By utilizing hierarchical clustering of the expressed transcripts, we identified a subset of known eye development genes expressed during early (st. 24 and st. 27) and late (st. 35) retinal development (Fig. 2A). For example, the transcription factor *myc,* which is expressed in retinal stem cells located in the ciliary marginal zone (CMZ) [35], was upregulated at st. 27 compared to st. 35 (Fig. 2A). A second cluster of development genes including *pitx3,* expressed during lens development [36], and *crx,* a transcriptional activator that regulates photoreceptor differentiation, were upregulated at st. 35 [37] (Fig. 2A). We expected that genes associated with an absence of eye phenotype would likely be expressed prior to eye differentiation, however our results do not suggest a clear pattern between mutant eye phenotypes and the temporal expression of each gene at the timepoints we assessed. These data indicate that regulatory eye genes likely have diverse temporal expressions across development, where their function may differ depending on the context – i.e. specification of optic tissues or the completion of retinogenesis.

**Figure 2:**
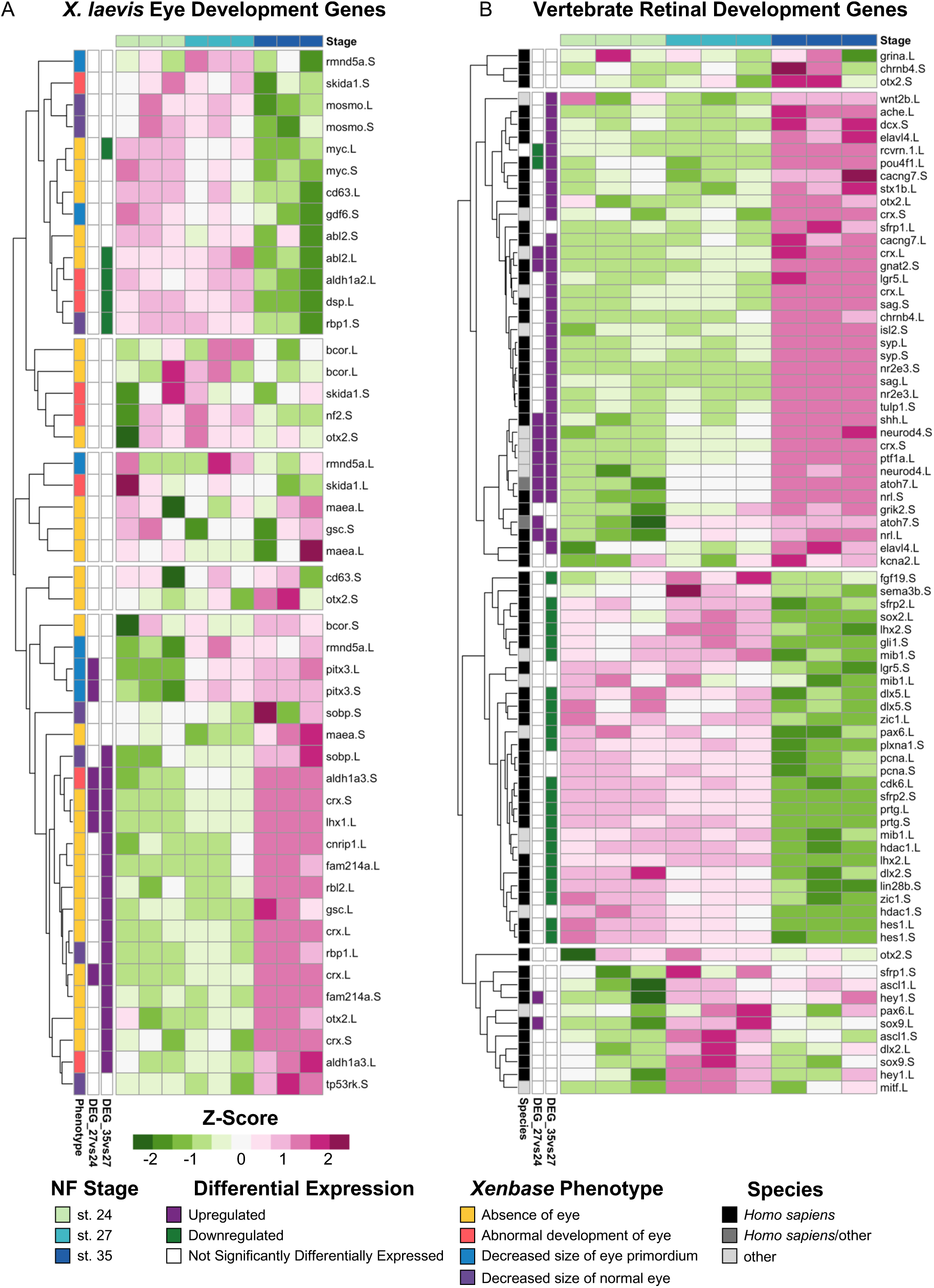
Expression of Known Vertebrate Eye Development Genes. (**A**) Relative expression (Z-score vst counts) of genes with known eye development phenotypes in *X. laevis* at st. 24, 27, and 35. The phenotype associated with each gene (Xenbase [32–34]) is displayed to the left. **(B)** Relative expression (Z-score vst counts) of vertebrate eye development genes (*H. sapiens*, *M. musculus*, and *D. rerio*) at st. 24, 27, and 35. Genes identified in the differential expression analysis (genes with adj. *p* < 0.05 and log_2_fold change ≥ 1 or ≤ –1) between consecutive developmental stages are displayed to the left (Purple: Upregulated, Green: Downregulated, White: Not differentially expressed).

Next, to investigate the extent to which transcriptional regulation of retinal development may be conserved across species, we analyzed the expression of genes belonging to six molecular superclusters previously identified from RNA-seq analysis of the developing human retina [38] (Fig. 2B). Two of these previously identified superclusters were associated with positive regulation of proliferation and neural development; one supercluster showed temporal upregulation of expression during early fetal retinogenesis (from embryonic days 52-57) and downregulated expression during later eye development stages [38]. Several genes belonging to these superclusters [38], including atonal homolog-7 (*atoh7*), fibroblast growth factor-19 (*fgf19)*, and cell cycle genes such as proliferating cell nuclear antigen (*pcna*) and cdk6 were also upregulated during early *Xenopus* retinal development between st. 24 and 27 in our own molecular analysis (Fig. 2B). The four other superclusters identified in the molecular analysis of human retina development showed upregulated levels of expression during late stages of retinogenesis (from embryonic day 125-136) [38]; these clusters were associated with biological processes such as synaptic transmission, phototransduction, and autophagy, among others [38]. Our results mirror this pattern, where expression of genes from these superclusters tended to have increased expression levels at st. 35 (Fig. 2B). For example, expression of the glutamate receptor *grik-2*, neural retina leucine zipper (*nrl*), acetylcholinesterase (*ache*) all had increased levels of expression at st. 35 compared to stgs. 24 and 27 (Fig. 2B), which is similar to the temporal expression pattern observed for human fetal retinogenesis [38]. Interestingly, genes in an autophagy-associated supercluster that were upregulated in late fetal retinogenesis in humans [38], such as *sema-3b* (semaphorin-3b) and *plxna-1* (plexin-A1), were downregulated in st. 35 *Xenopus* eyes (Fig. 2B). Collectively, our analysis of genes associated with human fetal retinal development suggests that the temporal expression of molecular signatures of early versus late retinogenesis may be conserved across vertebrate species.

Although there is heterogeneity in the RPC population at the same development stage, vertebrate RPCs have been classified into two groups based on conserved vertebrate transcriptional states identified from scRNA-seq studies [39]. “Primary RPCs” show increased expression of cell cycle regulatory transcripts and are largely proliferative whereas “neurogenic RPCs” express neural genes and undergo asymmetric cell division where one daughter cell differentiates into a retinal cell [39]. We examined conserved vertebrate gene markers that are characteristic of primary (proliferative) RPCs and neurogenic (differentiating) RPCs (Supplemental Table 3). Differentially expressed (adj. *p* ≤ 0.05, log_2_fold change ≥ 0.5) primary RPC markers were downregulated at st. 35 compared to st. 27. For example, proliferative RPC marker *sfrp2.L* was significantly downregulated (log_2_fold change = –2.41) from st. 27 to st. 35 but was not differentially expressed between st. 24 and 27 (Supplemental Table 3). Conversely, 8/9 differentially expressed (adj. *p* < 0.05, log_2_fold change ≥ 0.5) neurogenic RPC markers including *rlbp1*, *car2*, and *crym* were upregulated at st. 35 compared to st. 27 (Supplemental Table 3). The only downregulated neurogenic RPC marker at st. 35 was *sox8.S* (log_2_fold change = –0.87), which was previously shown to be expressed in cycling RPCs that give rise to Müller glia [40]. These data consistently show that RPCs in developing *X. laevis* eyes express conserved RPC markers in the temporal pattern observed in other vertebrate species, where primary RPC markers are more highly expressed during early retinogenesis (st. 24 and st. 27) and differentiating, neurogenic RPC markers are expressed during late retinogenesis (st. 35).

### Gene Ontology (GO) Analysis of Developing Optic Tissues

To further explore temporal differences in the molecular profiles of the developing optic tissues, we utilized Gene Ontology (GO) analysis [41] to uncover enriched biological processes and molecular functions in DEGs identified between the three developmental stages (Fig. 3). Biological processes involved in tissue development were enriched during early retinogenesis (st. 24 and st. 27) whereas GO terms associated with neuronal function and sensory perception were enriched during the later stage of eye formation (st. 35) (Fig. 3A). For example, we detected upregulation of ‘retina development in camera-type eye’, ‘nervous system development’, and ‘cellular developmental process’ at st. 27, whereas these biological processes were not enriched at st. 35 (Fig. 3A). ‘Neuron differentiation,’ ‘visual perception,’ and ‘neurotransmitter transport’, were enriched at st. 35, in addition to several metabolic/biosynthetic processes (including lipid metabolism and ATP biosynthesis) (Fig. 3A).

**Figure 3:**
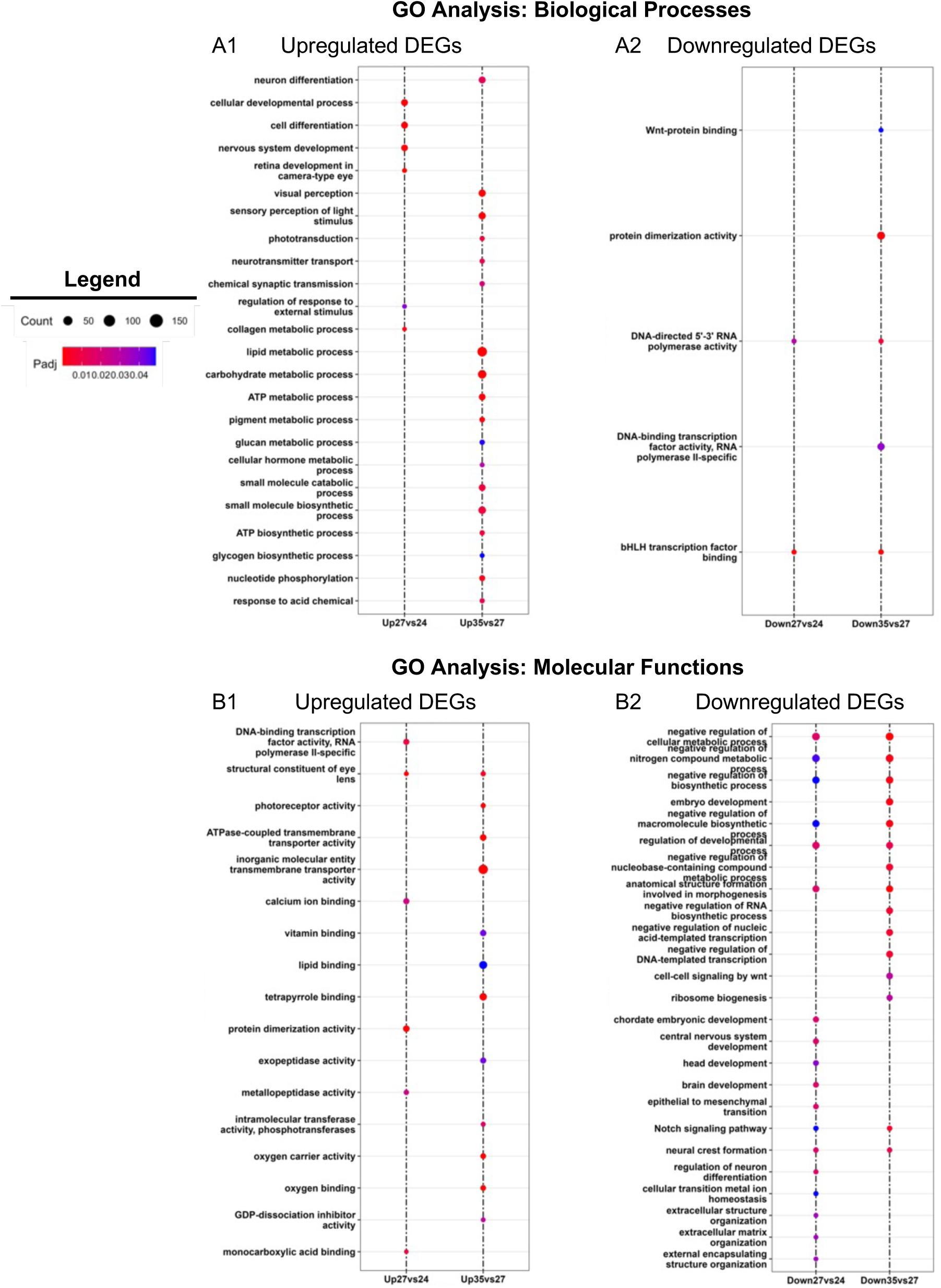
GO Enrichment Analysis. Gene Ontology (GO) analysis of statistically significantly differentially expressed genes (|log_2_ fold change| ≥ 1 and adj. *p* < 0.05) between consecutive developmental stages (st. 27 v 24 and st. 35 v 27). Enriched GOs for **(A)** biological processes and **(B)** molecular functions.

We detected similar patterns in molecular functions at each stage as well, where trends in GO term enrichment follow a pattern where eye development molecular programs are upregulated during early retinogenesis while the transcriptional landscape begins moving towards upregulation of metabolic processes and neuronal activity associated with acquisition of visual function during late retinogenesis (Fig. 3B). For instance, GO terms like ‘head development’ and ‘brain development’ were downregulated at st. 27 compared to st. 24 (Fig. 3B). Furthermore, upregulation of ‘photoreceptor activity’ and ‘calcium ion binding’ were GO terms more highly enriched at st. 35 versus st. 27 (Fig. 3B). Collectively, our analyses support previous work indicating *X. laevis* eyes are largely differentiated by st. 35 and gain some initial function during the late tailbud-free swimming stages [42,43]. Moreover, the data suggests that st. 27 is a dynamic period in eye development, where molecular functions shift from RPC proliferation and retinogenesis to transcriptional programs associated with differentiation of mature, functioning neurons.

### KEGG Pathway Analyses Identify Hedgehog, PPAR, and Wnt Signaling as Potential Regulators of Eye Development

Similar to GO analysis, we conducted Kyoto Encyclopedia of Genes and Genomes (KEGG) pathway analysis on DEGs (log_2_ fold change ≥ 1 and adj. *p* < 0.05) upregulated at both stages 24 and 27 during RPC proliferation and DEGs upregulated at st. 35 (Fig. 4 and Fig. S1) to highlight molecular pathways that regulate retinogenesis. Consistent with the timing of active RPC proliferation, over 60 cell-cycle related genes were upregulated at the two early timepoints (st. 24 and 27). Our KEGG analyses indicated that canonical developmental pathways including Wnt and Hedgehog signaling were also upregulated at early timepoints. This observation is consistent with previous studies indicating that Wnt and Hedgehog signaling can regulate neural precursor proliferation [44–46] (Fig. 4A). Hedgehog signaling is required for eye development in several models [52]. It has also been studied in the context of differentiation and patterning of the eye, indicative of its multiple roles during retinogenesis (Fig. 4A). Similarly, both canonical and non-canonical Wnt signaling are well established regulators of vertebrate eye development and mutations in pathway components result in eye abnormalities [47,48]. However, the requirement for signaling during different stages of eye development appears to be highly complex and species-dependent [48]. In *X. laevis*, *fzd-5* is required for RPC proliferation and retina development [49], but neurogenesis in the developing mouse retina is not dependent on *fzd-5* [50]. Given that Wnt signaling was also enriched at st. 27, it may play a role in regenerative neurogenesis as well. The role for Wnt signaling in embryonic eye regrowth is an exciting direction for future studies as it is a highly conserved regeneration signaling pathway [51].

**Figure 4:**
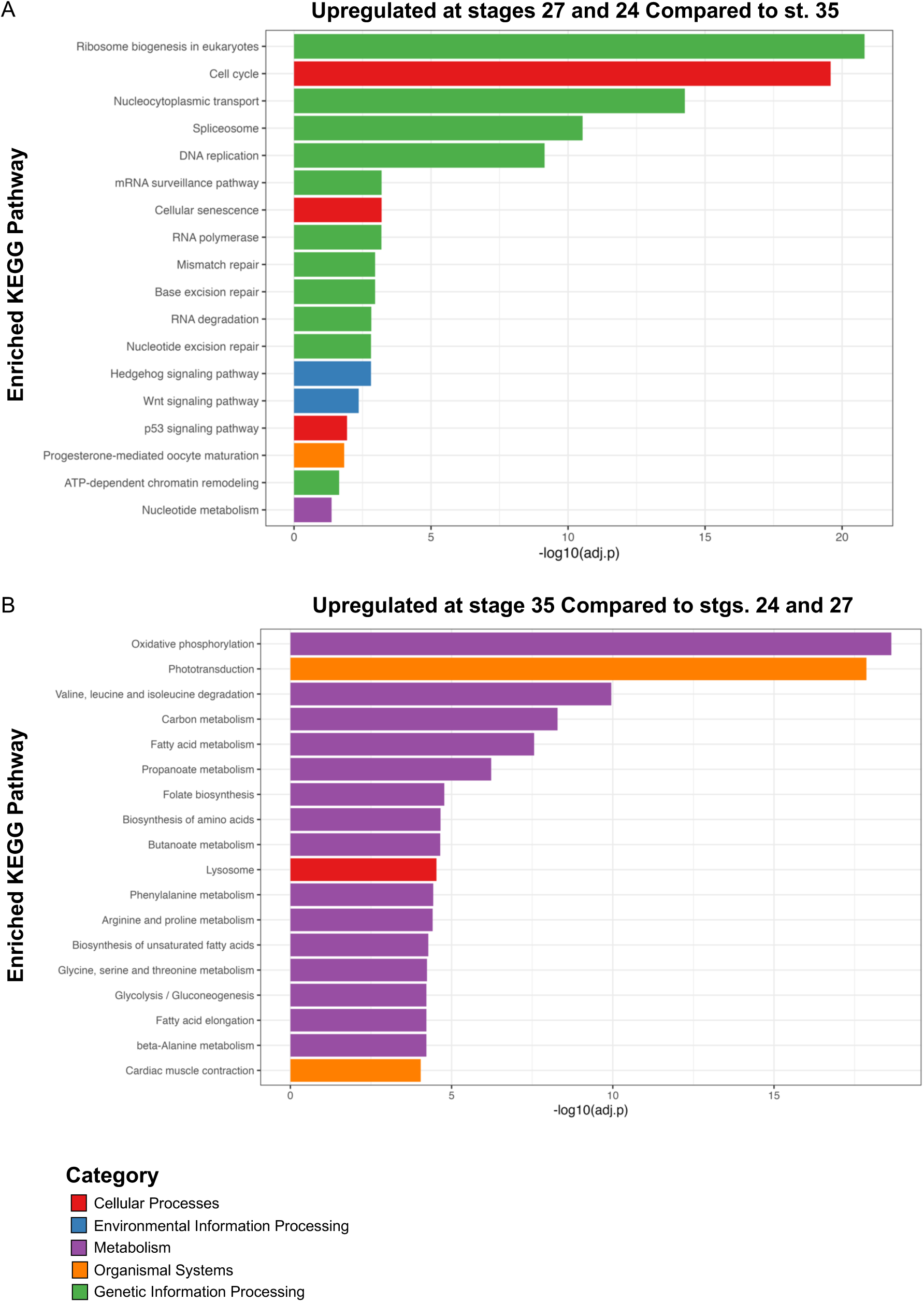
KEGG Pathway Enrichment Analysis. Kyoto Encyclopedia of Genes and Genomes (KEGG) analysis of **(A)** statistically significantly upregulated genes at st. 24 and 27 compared to st. 35 and **(B)** statistically significantly upregulated genes at st. 35 compared to st. 24 and 27. Only top hits are shown for (B) for full list of KEGG pathways see Supplemental Figure 1.

As opposed to st. 24 and 27, we detected enrichment for a variety of metabolic and biosynthetic pathways in differentiating st. 35 optic tissues. Pathways associated with neuronal function such as “Phototransduction” and “Cardiac muscle contraction” were also highly enriched (Fig. 4B). The full list of enriched pathways is shown in Fig. S1. The mature retina is highly metabolically active [53], thus it was not unexpected that a number biosynthetic pathways were upregulated during retinal differentiation. However, the spatiotemporal regulation of metabolism and molecular biosynthesis has not been well-investigated in developing *Xenopus* nervous tissues. It was previously shown that a glycolysis gradient arises in the developing chick neural tube in response to establishment of an FGF/Wnt signaling gradient; the graded glycolytic activity is required for axis elongation and cell fate specification in the tail bud [54]. Similarly, metabolic reprogramming during embryogenesis has been suggested as a mechanism to control stem cell differentiation and epigenetic state [55]. Our data suggest that metabolic reprogramming may occur during the late phase of retinal development, but whether this occurs in parallel with the onset of neuronal function or is associated with changes in differentiation of RPCs and RPC progeny remains unclear and is a potential area of future investigation. Given that optic tissues have high metabolic demands [53] and that we detected enrichment for phototransductive pathways (Fig. 4B), our KEGG analysis further suggests a switch to molecular programs associated with neuronal function during late retinal development.

### Conserved Eye Regeneration Gene Expression Analysis Across Retinal Development

Our recent work has shown that *X. laevis* eyes ablated at st. 27 are regrowth-competent and restored to age-appropriate size by 3 days post-surgery, where cell types are reborn in the order that parallels developmental retinogenesis [8,31,42]. Significantly increased levels of RPC proliferation during the first twenty-four hours post injury facilitated regrowth and led to a temporal shift in retinogenesis not observed during normal development [8]. The developing eye transcriptome generated here temporally overlaps with regrowth-competent embryonic stages (from st. 24 to st. 28), where our differential expression and GO analyses indicate that gene expression in optic tissues at st. 27 is highly dynamic. Plasticity in gene expression at this time may help to explain eye regrowth competency at this particular developmental stage. Given that the embryonic eye regrowth model was recently established and distinct from well-studied models of adult eye regeneration in other model systems such as *Danio rerio*, *Ambystoma mexicanum*, and *Schmidtea mediterranea,* we asked if the *Xenopus* eye developmental transcriptional program has components shared with adult eye regeneration processes. We compiled a representative list of genes that are required for eye regrowth in these model systems and queried their expression in our dataset. Of the regeneration genes identified from the literature, most had increased expression levels at st. 24 and 27, whereas expression levels were reduced at st. 35 (Fig. 5). However, a smaller subset of genes including the eye regeneration-associated orthodenticle homeobox transcription factors [56] (*otx-2*) and matrix metallopeptidase-9 (*mmp-9*) [57], had increased expression levels at st. 35 as compared to st. 24 and 27 (Fig. 5), suggesting they are unlikely to play a role in embryonic eye regrowth.

**Figure 5:**
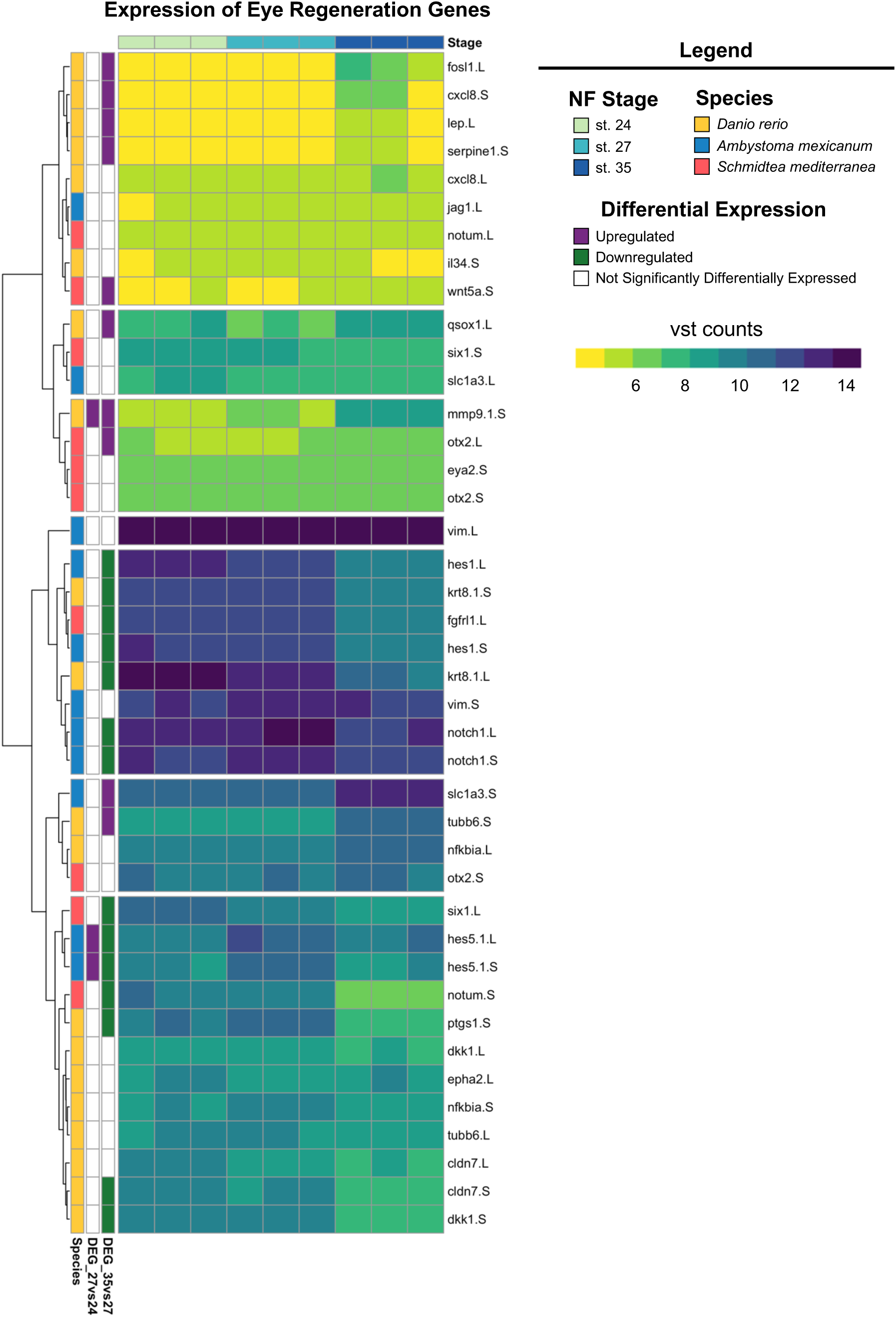
Expression of Known Eye Regeneration Genes. Expression (vst counts) of known eye regeneration genes in vertebrates (*D. rerio* and *A. mexicanum*) and the invertebrate *S. mediterranea* at st. 24, 27, and 35. Genes identified in the differential expression analysis (genes with adj. *p* < 0.05 and log_2_fold change ≥ 1 or ≤ –1) between consecutive developmental stages are displayed to the left (Purple: Upregulated, Green: Downregulated, White: Not differentially expressed).

The Notch signaling pathway is highly conserved across metazoans and plays essential roles in regulating proliferation, specification of cell fate, and coordinating differentiation during embryonic development and during adult tissue homeostasis [58–60]. Regeneration of diverse tissue types (including heart, liver, bone, and retina [61–64]) in many animal models also requires Notch signaling. Interestingly, genes encoding core canonical Notch signaling components, including the receptor *notch-1* and target gene *hes-5* were significantly upregulated at st. 27 compared to st. 24 or 35 (Fig. 5), which was consistent with our KEGG pathway analysis showing enrichment of Notch signaling specifically at st. 27 (adj. *p* = 1.75×10^−16^). Importantly, our recent study showed that *notch-1* is required for RPC proliferation and embryonic eye regrowth in *X. laevis*, but was not required for differentiation of regrowing optic tissues [65], supporting the analyses shown here. However, the potential mechanisms by which Notch signaling regulates embryonic eye regrowth remain unknown.

Using our transcriptional analyses, we identified genes belonging to the canonical Notch pathway, known target genes of Notch signaling, and non-canonical Notch pathway components and visualized their relative expression across eye development at st. 24, 27, and 35 (Fig. 6). Hierarchical clustering revealed that temporal expression of noncanonical and canonical Notch pathway components may differ, where many target genes and canonical signaling components were significantly upregulated at st. 27 such as: *hes5*, *dll1*, *notch.1* and *notch.3*, *adam.17*, and *dlc* (Fig. 6). This is similar to findings in mammalian models and zebrafish where cycling RPCs also show expression of several *hes/her* genes [39,66–68]. Interestingly, many genes associated with noncanonical Notch signaling such as Wnt ligands and SMAD genes had higher relative expression at either st. 24 or st. 35 compared to st. 27 (Fig. 6). Our data suggest further investigation into the role of Notch signaling in both retinogenesis and embryonic eye regrowth may reveal molecular drivers of RPC proliferation.

**Figure 6:**
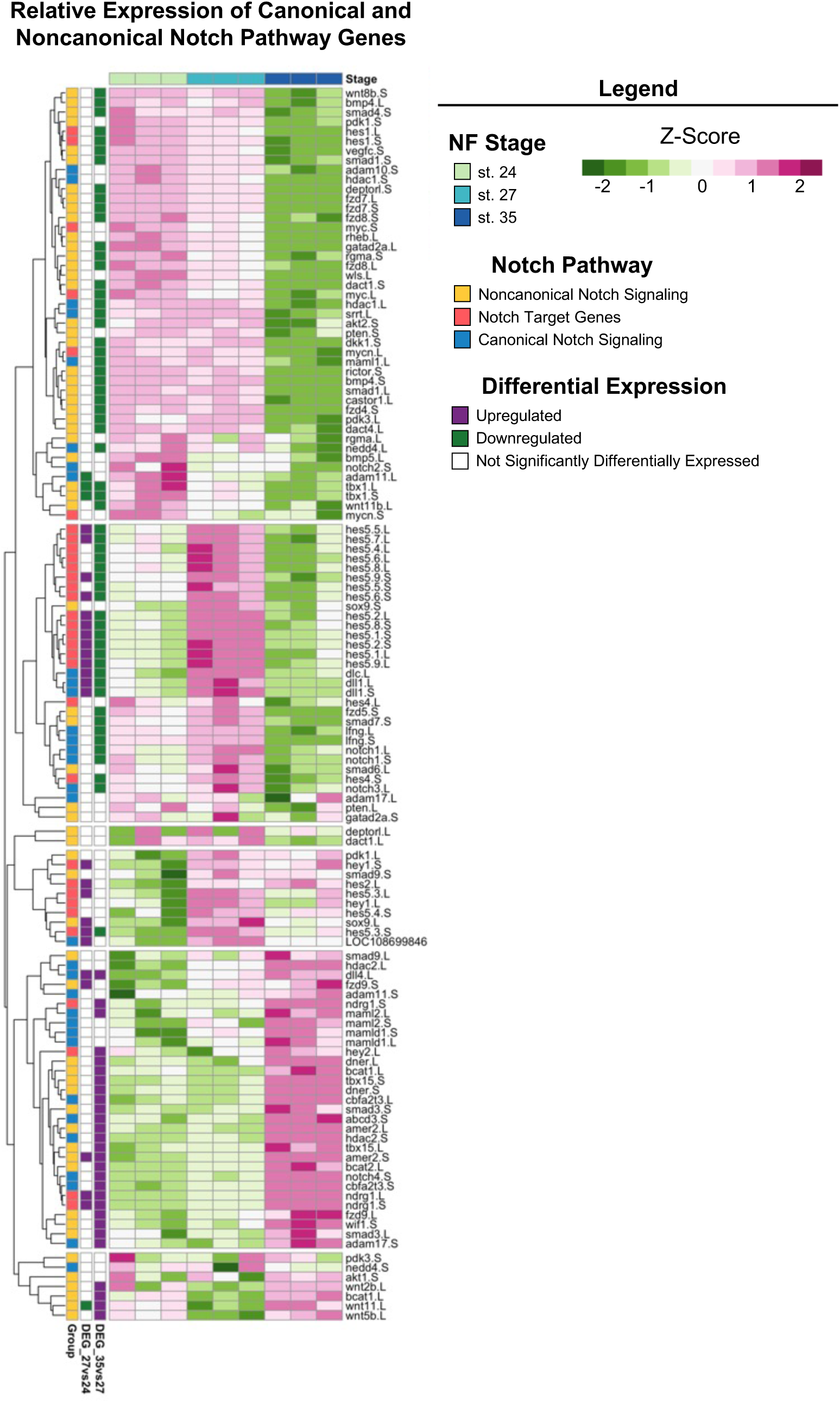
Notch Signaling Across Retinal Development. Relative expression (Z-score vst counts) of known Notch pathway genes (including genes that participate in canonical and noncanonical signaling, as well as downstream target genes) at st. 24, 27, and 35. Rows are ordered by hierarchical clustering and columns are ordered by stage. Genes identified in the differential expression analysis (genes with adj. *p* < 0.05 and log_2_fold change ≥ 1 or ≤ –1) between consecutive developmental stages are displayed to the left (Purple: Upregulated, Green: Downregulated, White: Not differentially expressed).

### Long versus Short Gene Homeologs are Differentially Expressed During Eye Development

Previous work has shown that homologous pairs of genes located in the L and S subgenomes of the allotetraploid frog *X. laevis* have correlated expression levels, where most genes show slight expression bias towards the L homeolog [69,70]. However, a small subset of genes has a strong expression bias, where the second copy (typically the S homeolog) has almost undetectable expression or has been lost from either subgenome entirely [70–72]. Given that L and S homeologs could be differentially expressed at unique developmental stages, we investigated whether retinal development follows the same patterns of subgenome expression observed in other studies.

We identified 3,489 L and S homeologs within our dataset; their Spearman correlation ranged between –0.97 and 1 and the Euclidean distance ranged between 0.25 and 38.25. Most of them can be considered similar across developmental time given their high correlation and minimal distance (Fig. 7A). Many genes from this subset, including *nrl*, *tbx1*, *sox-3*, *jund*, and *crybb1* have known roles in eye development, or embryogenesis more broadly [73–77]. A subset of genes also displayed extremely high distribution in relative expression, where the L variant was preferentially expressed in 1,284 of 3,489 gene pairs (Fig. 7B). Interestingly, some genes identified as having high distribution between L and S homeologs were categorized as ribosomal proteins including *rps4, rps5, rps12, rps15, rpl6, rpl11, and eif3c* (Supplemental Table 4). Other groups of genes that displayed high variance between subgenome homeologs included chaperone proteins (*cct2* and *cct7*), genes associated with proliferation (*azin1*, *smc3*, *hmgb2*, *and ybx1*), and members of protein kinase C signaling (*rack1* and *kpna2*) (Supplemental Table 4).

**Figure 7:**
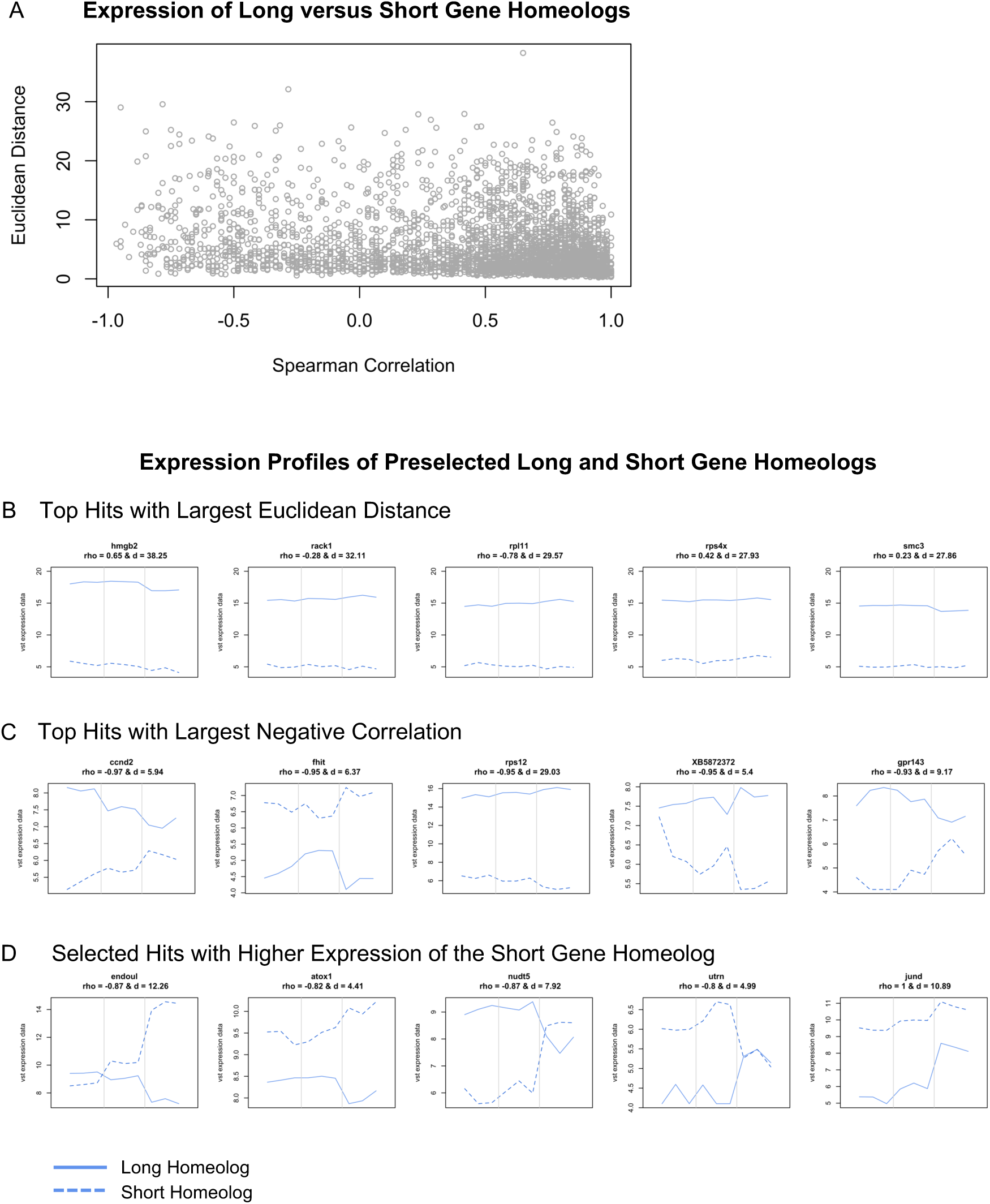
Expression Analysis of Long and Short Gene Homeologs Across Retinal Development. (**A**) Scatterplot of Spearman’s correlation coefficient versus the Euclidean distance of L and S homeologs. **(B-D)** Expression profiles (vst counts) of selected hits identified in (A). Homeologs with (B) largest Euclidean distance, thus with largest expression difference over developmental stages, (C) highest negative correlation, thus behaving differently over developmental stages, and (D) short gene homeologs with higher absolute expression than their long homeolog at some developmental stages. Dashed lines: Expression of the short homeolog, Solid lines: Expression of the long homeolog. The Spearman’s correlation coefficient (rho) and Euclidean distance (d) for homeologs are displayed at the top of the corresponding expression profile.

Lastly, we identified several genes where the expression of L and S homeologs was negatively correlated across development (Top hits shown in Fig. 7C), and many of these displayed preferential expression of the S homeolog at either st. 27 or st. 35. These included *endoul*, *atox1*, *nudt-5*, *ccnd-2,* and *uturn* (Fig. 7D). For example, expression of the long and short homeologs of *endoul*, also known as placental protein 11 (*pp11*), were similar at st. 24 and 27 but *endoul.S* expression was highly upregulated at st. 35 (Fig. 7D). Likewise, *nudt5.S* had elevated expression levels compared to the *nudt5.L* at st. 35 (Fig. 7D). Other genes like *atox1.S* and *utrn.S* had higher expression levels at all stages compared to the long homeolog of each gene (Fig. 7D), where *utrn.S* had upregulated expression specifically at st. 27 (Fig. 7D). Identification of significant differences in transcriptional profiles of developmentally relevant genes including *endoul* suggests that L and S subgenomes may differentially contribute to different retinal developmental processes. While subgenome-specific mutations may have arisen in many genes it is currently unclear why some homeologs are preferentially expressed when their sequence or protein domains are conserved between subgenomes. Given the L subgenome is typically used for analysis of RNA-seq datasets, our findings highlight that analysis of both L and S homeologs is an important step for identifying candidate genes during differential expression analysis. Lastly, it is now appreciated that the L subgenome developed after the S subgenome, and that many genes have been lost from S [69]. It will be interesting to consider how evolution of the allotetraploid subgenomes may have driven development of regenerative abilities or species-specific metamorphic processes, or whether retention of expression from S homeologs confers advantages during distinct developmental stages.

## Conclusions

A first step towards generating therapeutics to treat ocular diseases is improving our understanding of the molecular drivers of eye development and successful repair. Here we used RNA-sequencing to define the temporal transcriptomes of the *X. laevis* eye across three distinct developmental stages that span the early retinal development period of active RPC proliferation (st. 24 and 27) and then retinogenesis (st. 35). Identification of DEGs during retinal developmental stages coupled with GO and KEGG enrichment analyses indicated that the eye transcriptome between st. 24 to st. 35 is highly dynamic. Importantly, the tissues used for analysis included RPCs that must proliferate and differentiate properly to form the neural retina and RPE. Thus, future single-cell studies coupled with this bulk transcriptional analysis may uncover drivers of RPC proliferation and markers of retinal stem cells. Furthermore, we show that st. 27 optic tissues, which are regrowth-competent, express known markers of adult eye regeneration observed in other species. Specifically, Notch signaling was enriched at a regrowth-competent stage, indicating this key pathway may regulate RPC proliferation and maintenance during both development and regeneration. We also uncovered unique and variable expression profiles between long and short gene homeologs during retinal development. Our data indicated that an examination of both subgenomes for bioinformatic analysis is crucial for identifying potential molecular differences between expression of long and short gene homeologs during biological processes. Lastly, this study shows that *X. laevis* RPC transcriptomes are consistent with RPC marker expression patterns observed in mammalian models. This essential knowledge, combined with the distinct ability of *Xenopus* RPCs to endogenously restore the embryonic eye after injury, can facilitate ongoing investigations for pinpointing potential eye repair strategies.

## Materials and Methods

### Animal Care and Surgeries

Adult *Xenopus laevis* were purchased from Nasco (Janesville, WI) and grown via approved protocols and guidelines (UNLV Institutional Animal Care and Use Committee). Embryos were obtained via *in vitro* fertilization and cultured in 0.1× Marc’s Modified Ringer (MMR: 1 mM MgSO_4_, 2.0 mM KCl, 2 mM CaCl_2_, 0.1 M NaCl, 5 mM HEPES, pH 7.8) medium [78]. Prior to eye ablation, st. 24, 27, and 35 embryos [79] were anesthetized with MS222 (MilliporeSigma, St. Louis, MO). Fine surgical forceps (Dumont No. 5) were used to perform ablation surgery as described previously [8,80]. For imaging, embryos were fixed in MEMFA (100 mM MOPS (pH 7.4), 2 mM EGTA, 1 mM MgSO4, and 3.7% formaldehyde) and washed with PBS+0.1% Triton. Images were acquired using a Zeiss V20 Stereomicroscope with an Axiocam MRc camera.

### Sample Collection and RNA Extraction

RNA samples were collected from *Xenopus laevis* embryos at three district developmental stages: 24, 27, and 35. Each stage was collected in triplicates (n = 3) and each biological replicate was composed of pooled eye tissues from 20-30 animals/embryos. Eye ablation and tissue dissociation was performed in Liberase (Thermolysin Medium Research Grade, Roche, Indianapolis, IN) and/or calcium-magnesium free Modified Ringer solution (CMF-MR) (adapted from [81]). Dissociated tissues were immediately stored in TRIzol Reagent (Invitrogen, Carlsbad, CA) at −80°C. Total RNA was isolated using the Direct-zol RNA Microprep Kit (Zymo Research) according to the manufacturer’s instructions.

### Library Preparation and Sequencing

RNA-seq libraries were prepared from the extracted RNA using the Illumina TruSeq Stranded mRNA Library Prep Kit, utilizing 0.8 to 3.2 µg of input RNA. A diluted “A-Tailing Control” was introduced during ligation. Library quality was assessed using Agilent Bioanalyzer and molarity of the libraries was quantified using Roche KAPA qPCR. Paired-end sequencing with 150 cycles was performed on an Illumina NextSeq500 platform, and raw sequencing data were obtained from BaseSpace.

### Data Preprocessing

Adapter contamination in the raw sequencing data was assessed and contamination levels below 0.1% were considered acceptable. Overrepresented sequences were identified and removed using Fastp v0.20. Pre-trimming reports were generated to assess pre– and post-trimming sequence quality.

### Reference Genome and Annotation

The *Xenopus laevis* reference genome (XENLA_10.1_genome.fa) and annotation file (XENLA_10.1_GCF_XBmodels.gff3) were downloaded from Xenbase. The GFF3 annotation file was filtered to convert “pseudo” features to “exons” without gene IDs using the ‘agat_sp_filter_feature_by_attribute_presence.pl’ script. The filtered GFF3 file was then converted to GTF format using ‘gffread’ to facilitate compatibility with alignment tools such as STAR and HISAT.

### Alignment, Quantification, and Quality Control & Assessment

Trimmed RNA-seq data were aligned to the *Xenopus laevis* reference genome using both HISAT2 v2.2.1 and STAR v2.7 alignment tools. FeatureCounts was utilized to quantify reads assigned to genomic features. Quality control reports summarizing alignment statistics, read trimming data, and feature assignment were generated for both HISAT2 and STAR alignments. Comparisons between HISAT2 and STAR alignments were performed to assess alignment efficiency, with STAR demonstrating superior performance in terms of reads assigned to features.

### RNAseq Data Analysis

FeatureCounts data derived from the STAR alignment were imported into the statistical software R, where all downstream analyses were conducted. First, featureCounts underwent a quality control assessment to ensure sample quality and well-quantified transcripts, including checking library sizes for consistency across all samples. Transcripts with at least 10 counts were considered well-defined. To remove potential noise, a low count filter was applied, removing transcripts that 1) had too many zero counts across the samples and 2) had too many counts less than 10, as these could be attributed to technical noise. Transcripts that were well-defined in at least one of the three experimental groups were retained in the analysis. DESeq2 was used to process the filtered counts, and the variance stabilization transformation (VST) data were used for all visualizations. Principal Component Analyses (PCA) were conducted on the raw, filtered, and transformed data to ensure data integrity. We conducted all pairwise comparisons to determine differential expression. Transcripts with a false discovery rate (FDR) corrected *p*-value of less than 0.05 and an absolute value of log2 fold change of greater than or equal to 1 were deemed statistically significant and were used for Gene Ontology (GO) enrichment analysis and KEGG pathway analysis.

In addition to differential analysis, we further investigated the long (.L) and short (.S) versions of all transcripts. All transcripts with a L and S version were extracted. For each pair (e.g., lmo3.L and lmo3.S), we calculated the Spearman correlation (*rho*) and Euclidean distance (*d*) of the VST data to determine gene pairs with similar behavior or expression across developmental stages, respectively. We focused on three groups: 1) different expression magnitude (*d* > 27), 1), 2) different expression patters (*rho* < –0.9), and 3) ‘identical’ pattern and expression (*rho* < –0.9 and d < 1). We also extracted gene pairs of particular interest given prior knowledge to determine if their L and S version demonstrated different expression.

### Enrichment Analysis

#### Data Acquisition and Differential Expression Analysis

Gene expression data at different developmental stages (24, 27, and 35) of *X. laevis* were analyzed to identify differentially expressed genes (DEGs). DEGs were determined based on a criteria of a log2 fold change threshold of 1 and an adjusted p-value of 0.05. Comparisons were made between consecutive developmental stages (27 vs 24 and 35 vs 27) to identify upregulated and downregulated genes.

#### Annotation and Enrichment Analysis

The Entrez IDs of the DEGs were annotated using the *X. laevis* annotation hub database (org.Xl.eg.db). Gene Ontology (GO) and Kyoto Encyclopedia of Genes and Genomes (KEGG) pathway enrichment analyses were conducted on the annotated DEGs using the enrichGO and enrichKEGG functions, respectively, from the clusterProfiler package in R. Both analyses applied a significance cutoff of 0.05. To refine the GO enrichment results and eliminate redundancy, the simplify function, also part of the clusterProfiler package, was utilized to remove redundant and higher-level GO IDs.

#### Data Visualization

All statistical analyses and enrichment analyses were performed in R. Figures summarizing the results of the differential expression analysis, as well as GO and KEGG enrichment analyses, were generated using the ggplot2 package in R.

## Acknowledgements

This study was supported by grants from the National Institutes of Health (GM146672, GM103440, and U54 GM104944). This work was also supported by a Western Michigan University Dissertation Completion Fellowship to S.J.H. We thank Cindy Kha for tissue dissections and RNA isolation, the Nevada Genomics Center (Paul Hartley) for sequencing the samples, and the Nevada Bioinformatics Center (RRID:SCR_017802) (Lucas Bishop, Jeremiah B. Reyes, Hans Vasquez-Gross, and Richard Tillett) for bioinformatics support. We also thank Wendy Beane for comments and the Xenbase team for help with annotations.

## Author contributions

Conceptualization, K.A.-S.T.; formal analysis, S.J.H., and J.P.; investigation, S.J.H., J.P., and K.A.-S.T; resources, J.P. and K.A.-S.T; data curation, J.P.; writing—original draft preparation, S.J.H.; writing—review and editing, S.J.H., J.P., and K.A.-S.T; visualization, J.P. and S.J.H.; supervision and project administration, K.A.-S.T.; funding acquisition, S.J.H., J.P., and K.A.-S.T. All authors have read and agreed to the published version of the manuscript.

## Corresponding authors

Correspondence and material requested should be sent to Kelly Tseng.

## Data availability

Raw sequencing data have been deposited to the Sequence Read Archive (SRA) as part of https://www.ncbi.nlm.nih.gov/ and are accessible under accession number for PRJNA1124041. Other relevant data supporting the key findings of this study are available within the article, supplemental information, public repositories, or from the corresponding author upon reasonable request,

**Supplemental Figure 1:**
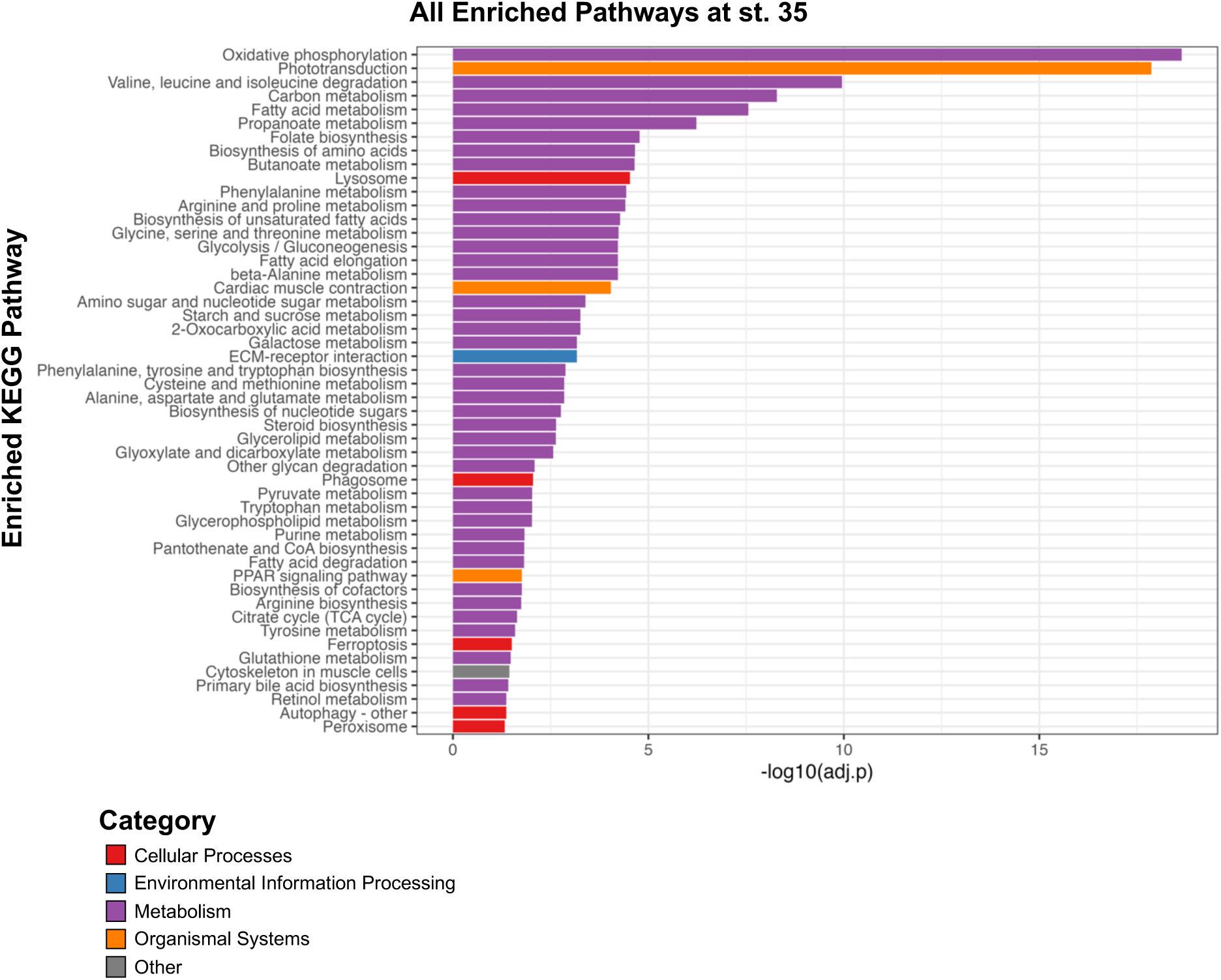
All Enriched KEGG Pathways at st. 35. All Enriched KEGG Pathways of statistically significantly upregulated genes at st. 35 compared to st. 24 and 27.

